# The interplay of stiffness and force anisotropies drive embryo elongation

**DOI:** 10.1101/095752

**Authors:** Thanh TK Vuong-Brender, Martine Ben Amar, Julien Pontabry, Michel Labouesse

**Affiliations:** Sorbonne Universités, UPMC Univ Paris 06, CNRS, Laboratoire de Biologie du Développement - Institut de Biologie Paris Seine (LBD - IBPS), 75005 Paris, France; Development and Stem Cells Program, IGBMC, CNRS (UMR7104), INSERM (U964), Université de Strasbourg, 1 rue Laurent Fries, BP10142, 67400 Illkirch, France; Laboratoire de Physique Statistique, Ecole Normale Supérieure, UPMC Université Pierre et Marie Curie, Université Paris Diderot, CNRS, 24 rue Lhomond, 75005 Paris, France; Institut Universitaire de Cancérologie, Faculté de Médecine, Université Pierre et Marie Curie-Paris 6, 91 Bd de l’Hôpital, 75013 Paris, France

## Abstract

The morphogenesis of tissues, like the deformation of an object, results from the interplay between their material properties and the mechanical forces exerted on them. Whereas the importance of mechanical forces in influencing cell behaviour is widely recognized, the importance of tissue material properties, in particular stiffness, has received much less attention. Using *C. elegans* as a model, we examine how both aspects contribute to embryonic elongation. Measuring the opening shape of the epidermal actin cortex after laser nano-ablation, we assess the spatiotemporal changes of actomyosin-dependent force and stiffness along the antero-posterior and dorso-ventral axis. Experimental data and analytical modelling show that myosin II-dependent force anisotropy within the lateral epidermis, and stiffness anisotropy within the fiber-reinforced dorso-ventral epidermis are critical to drive embryonic elongation. Together, our results establish a quantitative link between cortical tension, material properties and morphogenesis of an entire embryo.

## Introduction

Morphogenesis and organ formation rely on force distribution and tissue material properties, which are often heterogeneous and evolve over time. Forces are generated through a group of relatively well-conserved molecular motors associated with the cytoskeleton, among which myosin II linked to actin filaments is the most prevalent during epithelial morphogenesis (Vicente-Manzanares, 2009). Myosin II spatial distribution and dynamics greatly influence morphogenetic processes (Levayer and Lecuit, 2012). In particular, the asymmetric distribution of the actomyosin network and its pulsatile behaviour define the direction of extension during *Drosophila* germband elongation (Bertet, 2004, Blankenship, 2006), *Drosophila* renal tubule formation (Saxena, 2014) or *Xenopus* mesoderm convergent extension (Shindo and Wallingford, 2014). Whereas the implication of mechanical forces has been intensively investigated (Zhang and Labouesse, 2012, Heisenberg and Bellaiche, 2013), much fewer studies have considered the impact of tissue material properties *in vivo*, except for their influence on cell behaviour *in vitro* (Kasza, 2007).

*C. elegans* embryonic elongation represents an attractive model for studying morphogenesis, as it offers single cell resolution and powerful genetic analysis. During its elongation, the embryo evolves from a lima-bean to a typical cylindrical shape with a four-fold increase in length, without cell migration, cell division, or a notable change in embryonic volume (Sulston, 1983, Priess and Hirsh, 1986) (figure 1a). This process requires the epidermal actomyosin cytoskeleton, which acts mostly in the lateral epidermis (also called seam cells), while the dorso-ventral (DV) epidermal cells may remain passive (supplementary SI1)(Wissmann, 1997, Wissmann, 1999, Shelton, 1999, Piekny, 2003, Diogon, 2007, Gally, 2009, Chan, 2015, Vuong-Brender, 2016). Indeed, the non-muscle myosin II is concentrated in seam cells; in addition short disorganized actin filaments, which favour actomyosin contractility, are present in seam cells, but not in the DV epidermis where they instead form parallel circumferential bundles (Figure 1b-d)(Gally, 2009, Priess and Hirsh, 1986). The actomyosin forces are thought to squeeze the embryo circumferentially, to thereby increase the hydrostatic pressure and promote embryo elongation in the antero-posterior (AP) direction (Priess and Hirsh, 1986) (figure 1e).

**Figure 1:**
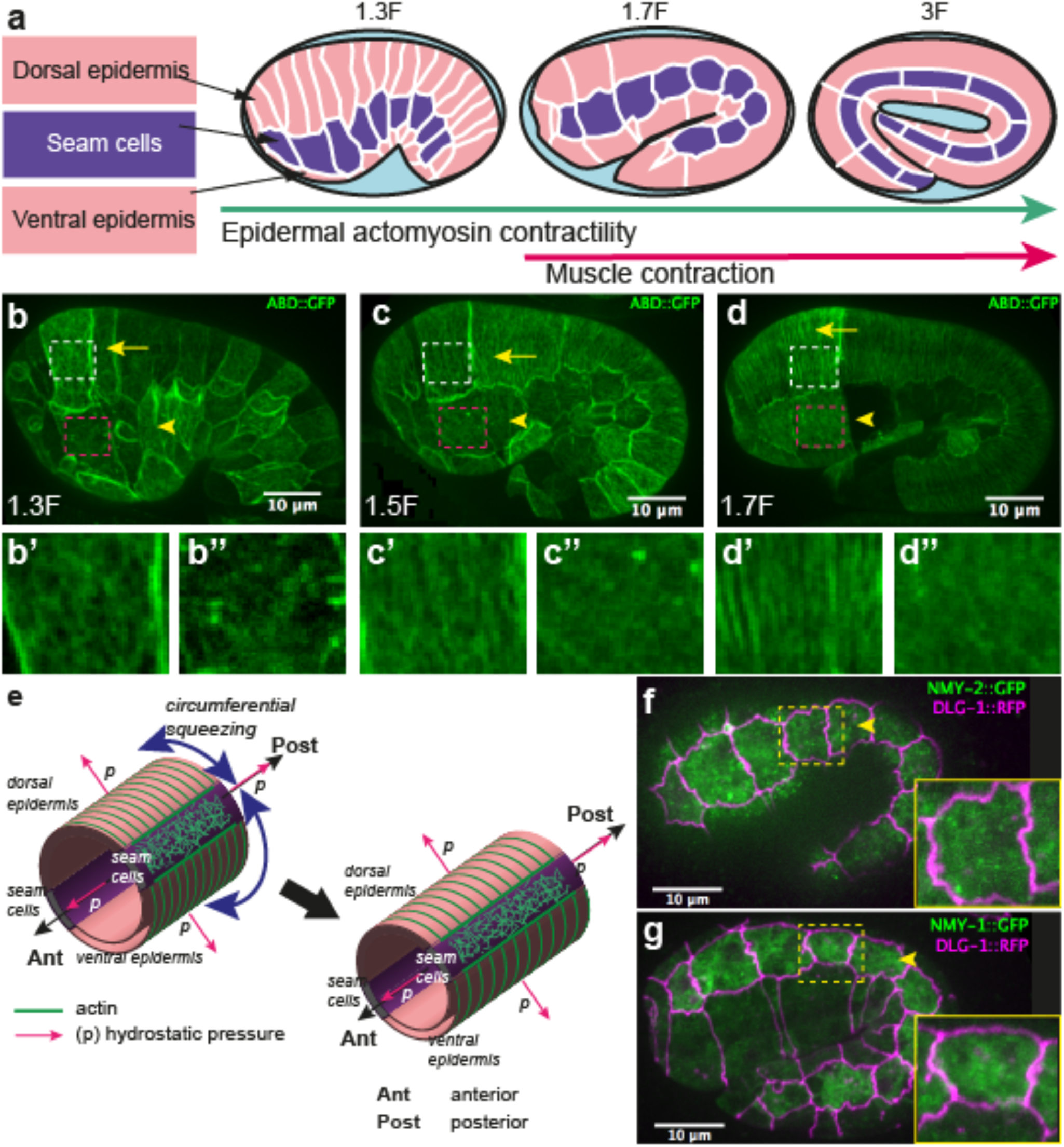
Overview of *C. elegans* embryonic elongation. (**a**) *C. elegans* embryonic elongation is driven in part by epidermal actomyosin contractility and in part by muscle contractions. The length of the embryo is used for staging: 2-fold (2F) stage means roughly 2-fold increase in length from the beginning of elongation. Representative stages are shown; anterior to the left, dorsal up. (**b, c, d**) Actin filament organization at the 1.3F, 1.5F and 1.7F stages, respectively, visualized with an ABD::GFP marker. Actin filaments progressively organize into circumferential parallel bundles in DV cells (arrows), arrowheads point to seam cells. Note that the integrated ABD::GFP marker shows some cell to cell variation of expression. (**b’, b”, c’, c”, d’, d”**) Zoom in of actin pattern in DV cells (rectangle above) and seam cells (rectangle below), respectively, of the images in (**b**), (**c**) and (**d**) respectively. (**e**) Actomyosin forces squeeze the embryo circumferentially to make it elongate in the antero-posterior direction. (**f, g**) Endogenous distribution of the two non-muscle myosin II isoforms visualized with CRISPR GFP-labelled myosin heavy chains NMY-1 and NMY-2, respectively. Arrowheads point to seam cells, which are delineated by the junctional marker DLG-1::RFP.

Although the published data clearly imply myosin II in driving elongation, they raise a number of issues. First, myosin II does not show a polarized distribution (figure 1 f-g) nor does it display dynamic pulsatile foci at this stage; hence, it is difficult to account for the circumferential squeezing. Moreover, force measurements are lacking to establish that the actomyosin network does squeeze the embryo circumferentially. Second, a mechanical continuum model is needed to understand how the embryo extends preferentially in the AP direction.

To address those issues, we used laser ablation to map the distribution of mechanical stress (i.e the force per unit area) and assess tissue stiffness (i.e. the extent to which it resists deformation) in the embryonic epidermis. We then correlated the global embryonic morphological changes with these physical parameters. Finally, we developed continuum mechanical models to account for the morphological changes. Altogether, our data and modelling highlight that the distribution of forces in the seam cells and the stiffness in the DV epidermis must be polarized along the circumferential axis (or DV axis) to drive elongation.

## Results

### Measuring the mechanical stress on the actin cortex through laser ablation

To measure the stress distribution on the actin cortex, we used laser nano-ablation, which has now become a standard method to assess forces exerted in cells, to sever the actin cytoskeleton and observe the shape of the opening hole (figure 2a). We visualized actin with a GFP- or mCherry-labelled actin-binding-domain protein (ABD) expressed in the epidermis (Gally, 2009) (figure 1b-d). We adjusted the region of interest to cut within one cell, restricting our analysis to the early phase of elongation (≤1.7F; for staging, see figure 1 legend).

**Figure 2:**
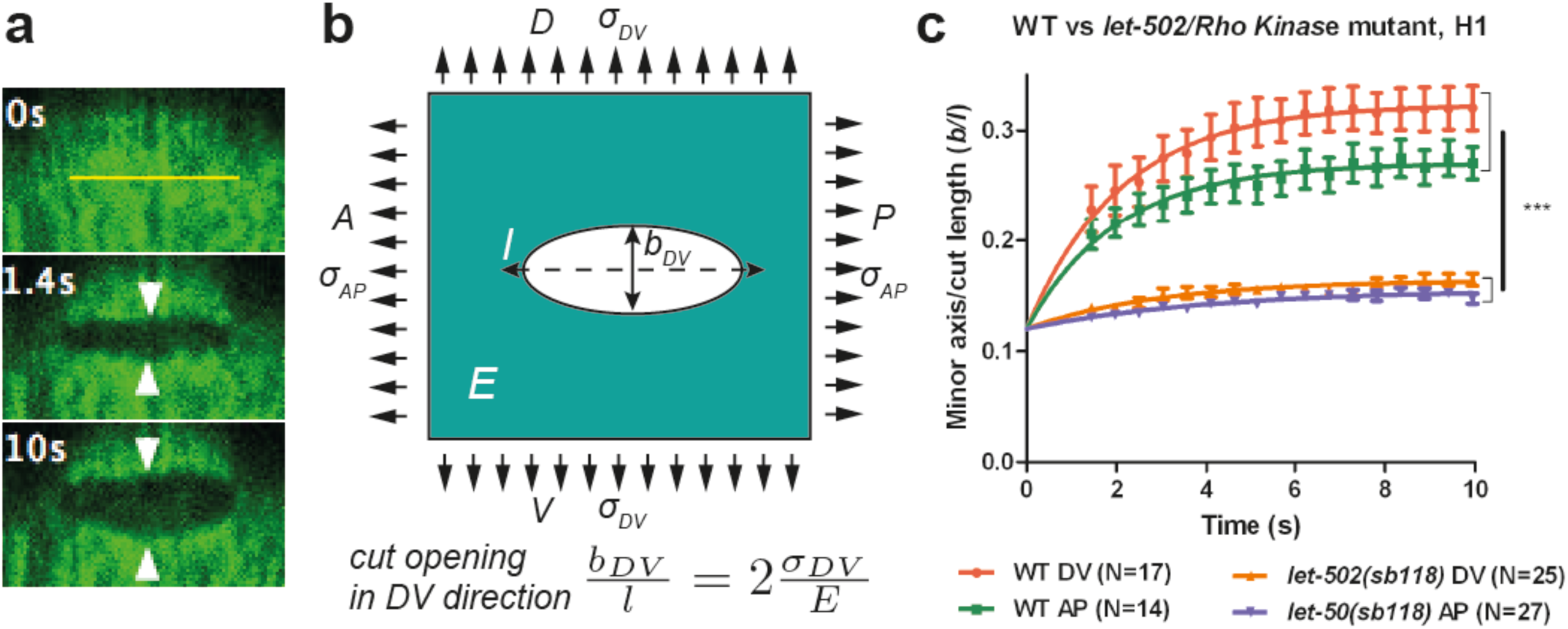
Physical model using the shape of the cut opening at equilibrium to measure the ratio of stress to Young modulus. (**a**) The GFP-labelled actin cortex of HYP7 dorsal epidermal cell at the 1.7F stage before (0 s), 1.4 s and 10 s after laser severing; cut along the AP direction with a 5 μm cut length. Double arrowheads, distance between cut borders, which increases with time. (**b**) Model of epidermal cells as an infinite elastic plane under biaxial stress in the AP and DV directions after an incision of length *l*. The final shape of the cut opening is an ellipse. The cut opening in the DV direction (after an incision along the AP direction) depends on the stress along the DV direction and the Young modulus (see text). (**c**) The opening depends on myosin II activity: comparison of the cut response in the seam cell H1 between wild-type (WT) and *let-502(sb118)/Rho kinase* mutant embryos at a stage when muscles start to twitch (around 1.5F). Time zero, moment of the cut; DV and AP show direction of opening. Two-tailed t-test, ***: p=4*10^−7^ between WT DV and *let-502(sb118)* DV, p=4*10^−6^ between WT AP and *let-502(sb118)* AP; N, number of embryos examined.

We observed two types of ablation responses (see Methods). In the first (accounting for > 80% of the cases), the opening hole within the actin cytoskeleton reached equilibrium in less than 10 s, and resealed within less than 2 min (Figure 2-Figure supplement 1, supplementary movie 1). In such embryos, actin occasionally accumulated around the cut borders but not around cell borders (supplementary movie 1). Imaging calcium levels, which can rise after laser wounding (Xu and Chisholm, 2011, Razzell, 2013, Antunes, 2013), showed either no change or a localized increase (supplementary movie 2, Figure 2 – Figure supplement 2a-b). In the second, an actin ring accumulated around the cell borders during the repair process (supplementary movie 3) and a calcium wave propagated to nearby epidermal cells (supplementary movie 4; Figure 2-Figure supplement 2c). Whereas embryos showing the first response continued to develop and hatched, those showing the second response arrested their development and eventually died. In all subsequent studies, we only took into account the first type of response, which should correspond to a local cortex disruption.

To compare the response between different conditions, we detected the cut opening shape, which we fitted with an ellipse to derive the shape parameters (see Methods). The laser setup we used did not enable us to image the recoil dynamics within the first second after the cut, which other investigators previously used to assess the extent of mechanical stress (Rauzi and Lenne, 2015, Smutny, 2015, Saha, 2016). To circumvent this issue, we developed a novel analysis method to derive mechanical stress, based on the equilibrium shape of a thin cut in an infinite elastic isotropic plane, subjected to biaxial loading (stress applied in two perpendicular directions)(Theocaris, 1986). The rationale for approximating the epidermis to such a plane is further outlined in the supplementary SI2-SI3. In these conditions, a thin cut will open to form an elliptical hole at equilibrium (figure 2a). The opening of the cut reflects mechanical stress in the direction perpendicular to the cut direction.

We cut the epidermal actin specifically in the AP and DV directions, which we found to correspond to the stress loading directions (figure 2b; supplementary SI2). For a cut in the AP direction, the minor axis of the ellipse at equilibrium, *b_DV_*, will be proportional to the cut length, *l*, and to the ratio of stress in the DV direction, *σ_DV_*, over the Young modulus *E* of the plane (Theocaris, 1986) (figure 2b):

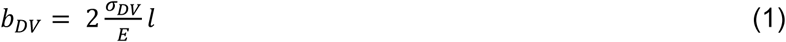

We will call the ratio
 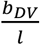 the opening in the DV direction of a cut made along the AP direction (figure 2b), and similarly 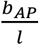 the opening in the AP direction of a cut made in the DV direction. Thus, we used the opening of the hole in a given direction to derive the stress in that direction.

We compared the conclusions drawn from this method with methods relying on the recoil dynamics (Rauzi and Lenne, 2015, Smutny, 2015, Saha, 2016) (supplementary SI3). The half-time of the cut border relaxation, which depends on the ratio of viscosity over stiffness, was similar in the AP and DV directions (supplementary SI3), supporting the hypothesis that the seam cell cortex is isotropic. We found an agreement between both methods for the AP versus DV stress ratio, and similar trends for the stress magnitude.

To further examine the validity of this method, we did two tests. First, the aforementioned theory (Theocaris, 1986) predicts that the minor to major axis ratio of the opening ellipse is independent of the initial cut length. We found that it is the case when the cut length varied from 3 μm to 6 μm (supplementary SI4). Second, to prove that the opening observed after laser cutting depends on myosin II activity, we performed cuts in embryos defective for the main myosin II regulator, LET-502/Rho-kinase (Diogon, 2007). As shown in figure 2c, the opening in the seam cell H1 at the 1.5F stage in *let-502(sb118ts)* embryos changed very little and was significantly smaller than in WT embryos, consistent with a decrease of mechanical stress.

Thus, we feel confident that the method based on the opening shape measures actomyosin-dependent stress and can reliably report on the differences of stress along the DV and AP directions.

### Stress anisotropy in seam cells correlates with embryonic morphological changes

We applied the method described above on three seam cells (head H1, body V3, tail V6; figure 3a), since myosin II acts mainly in seam cells (Gally, 2009), and compared the response with embryonic morphological changes. We focused on the anisotropy of stress between the DV and AP directions (difference of stress along both directions) in a given cell (figure 3b). Indeed, in other systems, such as *Drosophila* embryos (Rauzi, 2008) and *C. elegans* zygotes (Mayer, 2010), this parameter is critical.

**Figure 3:**
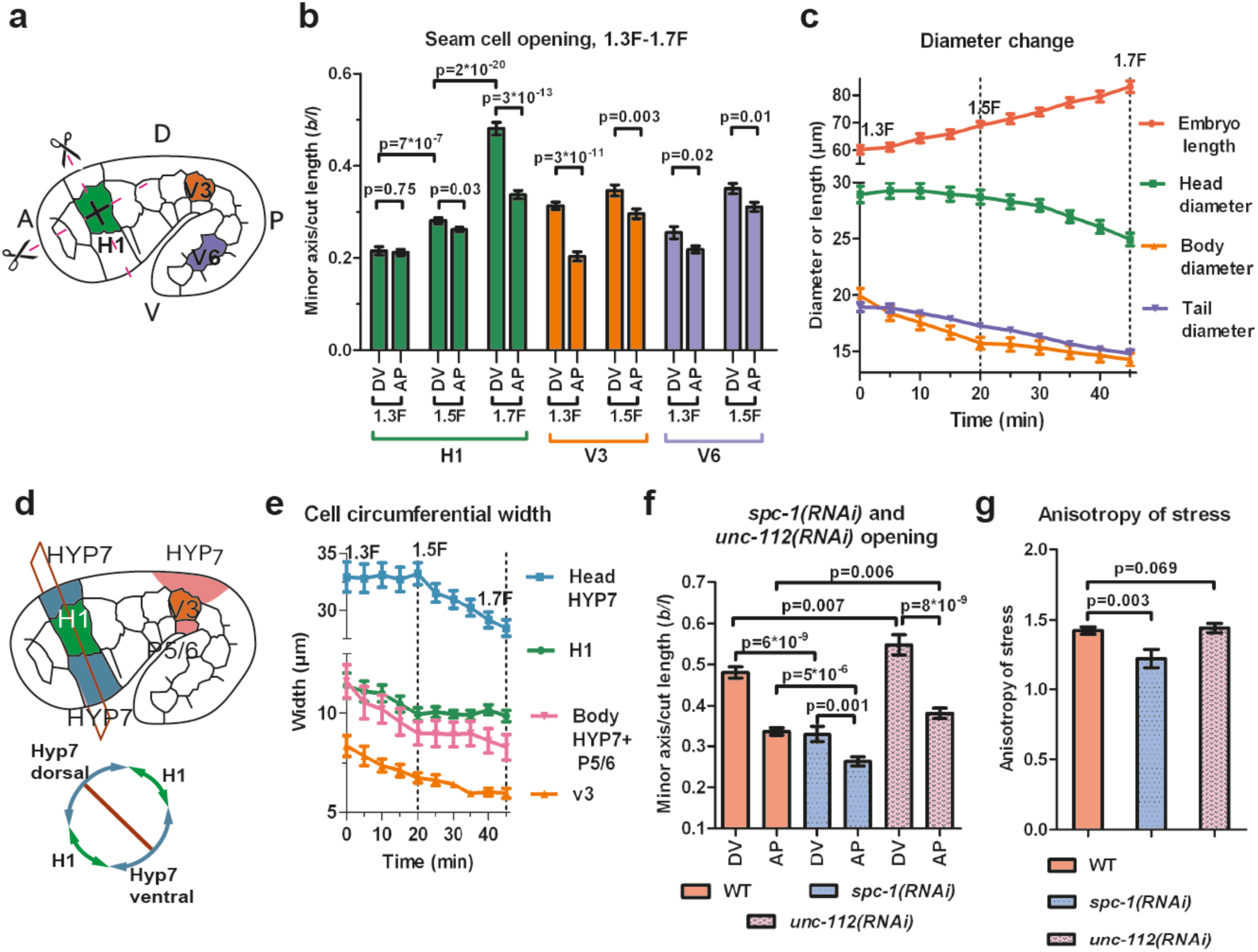
Stress anisotropy in seam cells correlates with morphological changes and partially depends on the spectrin cytoskeleton. (**a**) Scheme showing laser ablation experiments in the AP and DV directions for H1, V3 and V6 seam cells at different stages. A, anterior; P, posterior; D, dorsal; V, ventral. (**b**) Cut opening in H1, V3 and V6 from the 1.3F to the 1.7F stages (see figure 2b). p-value of two-tailed t-test is reported. (**c**) Changes in embryo length, head diameter at the level of H1, body diameter at the level of V3 and tail diameter at the level of V6, between the 1.3F and 1.7F stages. N=10. (**d**) Scheme showing the measurement of circumferential cell width in the head (above) and corresponding section (below). (**e**) The circumferential width of H1, V3, head and body DV cells (averaged for dorsal and ventral cells) are reported. N=10. (**f**) Measures of the cut opening in H1 for WT, *spc-1(RNAi)* treated and *unc-112(RNAi)* muscle defective embryos at a stage equivalent to the 1.7F stage. p-value of two-tailed t-test is reported. (**g**) Comparison of the stress anisotropy in H1, defined by DV/AP stress, between WT 1.7F stage, *spc-1(RNAi)* embryos and *unc-112(RNAi)* embryos at the equivalent 1.7F stage. p-value of Z-test is reported. The number of embryos used for ablation is given in supplementary table 1.

At the 1.3F stage in H1, there was no significant stress anisotropy; however, as the embryo elongated to the 1.5F and 1.7F stages, the stress became anisotropic (figure 3b). In V3, the anisotropy of stress evolved in the opposite direction, with higher stress anisotropy at the 1.3F compared to the 1.5F stage (figure 3b). In V6, the stress was slightly anisotropic at both the 1.3F and 1.5F stages (figure 3b). In all cells, whenever the stress became anisotropic it was higher in the DV direction. Overall, the opening increased as the embryo elongated from the 1.3F to the 1.5F stage and from the 1.5F to the 1.7F stage for H1.

To correlate the stress anisotropy with the morphological changes of the embryo, we used markers labelling cortical actin (an ABD) and junctions (HMR-1/E-cadherin). We observed that the head, body and tail diameter (at the level of H1, V3 and V6, respectively) decreased at different rates over time (figure 3c), as also observed by Martin and colleagues (Martin, 2014). The head diameter did not diminish between the 1.3F and 1.5F stages when the stress was nearly isotropic, but decreased significantly between the 1.5F and 1.7F stages as the stress anisotropy increased. Conversely, the body diameter decreased the fastest between the 1.3F and 1.5F stages when the stress was highly anisotropic, then changed at a lower pace beyond the 1.5F stage when the stress became less anisotropic. Finally, the tail diameter decreased nearly linearly between the 1.3F and 1.7F stages, at a lower rate than the body diameter, coinciding with a smaller anisotropic stress in V6. Thus, the local morphological changes within the embryo correlate with locally higher stress in the DV compared to the AP direction.

To define whether all cells equally contribute to the diameter change, we quantified the circumferential width of the epidermal cells H1, V3 and their adjacent DV cells (figure 3d-e). At the level of V3, the decrease in body diameter came from both seam (V3) and DV cells, whereas in the head it came mainly from DV cells (figure 3e). Collectively, our results strongly suggest that the stress anisotropy correlates with morphological changes. Furthermore, we found that both seam and DV epidermal cells contribute to the changes in embryo diameter, irrespective of their level of active myosin II.

### The establishment of stress anisotropy depends in part on the spectrin cytoskeleton

Taking the H1 cell as an example, we considered some cellular factors that could contribute to the stress anisotropy in seam cells: (i) actin-anchoring proteins, (ii) muscle-induced tension. To ease comparisons, we defined the anisotropy of stress (AS) as

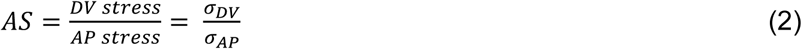

which can be derived from the ratio of the opening along the DV and AP directions, see equation (1).

First, we examined the actin-anchoring spectrin cytoskeleton, which is essential for embryonic elongation (Moorthy, 2000, Norman and Moerman, 2002, Praitis, 2005). In *spc-1(RNAi)* embryos, at a developmental timing equivalent to the 1.7F stage in control embryos, we found a smaller opening in both AP and DV directions and a decrease of AS compared to WT (figure 3f-g). This may account for the slower elongation rate of *spc-1(RNAi)* embryos and their arrest at the 2F stage (Figure 3-Figure supplement 1). Thus, spectrin partially contributes to the AS at the 1.7F stage.

Second, we wondered whether muscle contractions, which start after the 1.5F stage, could account for AS changes (figure 3b). Compared to controls, embryos depleted of UNC-112/Kindlin, which mediates sarcomere assembly (Rogalski, 2000), showed a significantly higher opening in both the AP and DV directions at the 1.7F stage, but no change in stress anisotropy (figure 3f-g). This is consistent with their wild-type elongation rate up to the 2F stage (Figure 3-Figure supplement 1). Thus, AS establishment in the H1 cell after the 1.5F stage is independent of muscle contractions.

### A mechanical model for seam cell elongation depending on stress anisotropy

To define the possible causal relationship between the AS and embryonic shape changes, we aimed at simplifying the shape of the embryo to apply classical physical laws such as the Young-Laplace equation, which predicts the relationship between surface tension and the surface curvature. As illustrated in figure 1, the embryo has a circular section and a cylindrical or conical shape depending on the stage, in which the epidermis is thin (100 nm to 2 μm, depending on areas; www.wormatlas.org) compared to the embryo diameter (25 μm). Within the embryo, the epidermis is subjected to hydrostatic pressure when the section decreases (Priess and Hirsh, 1986). We can thus model the *C. elegans* embryo as an isotropic thin-wall (the epidermis) vessel with capped ends under hydrostatic pressure, and determine the relationship between the mechanical stress on the epidermis and the embryo shape.

First, we calculated the anisotropy of stress on the wall of such a vessel. For an axisymmetric vessel, the AS on the wall depends on the surface curvature and the radius (supplementary SI5), which for simple geometrical configurations can be written as shown in figures 4a-c. Typically, the AS factor, or the DV to AP stress ratio, is equal to 1 for a sphere, equal to 2 for a cylinder and takes an intermediate value between 1 and 2 for an ellipsoid. We can simplify the geometry of *C. elegans* embryos as a curved cylinder (body), attached to a sphere (head) between the 1.3F and 1.5F stages (figure 4d-e). The head evolves into an ellipsoid between the 1.5F and 1.7F stages (figure 4f). Thus, the AS of the head can be determined easily. We previously observed that the AP stress among the seam cells at a given stage differs by 20% (figure 3b). Thus, if we approximate the AP stress as a constant at a given stage, the AS in the body will depend on the ratio of the body to head radius (figure 4d-e, supplementary SI5). Given the head and body diameter of the embryo (figure 3c), we can compare the AS predicted by the thin-wall vessel model with those derived experimentally with laser ablation (figure 4g). Both values are nearly identical, showing that the AS can be predicted based on the embryonic geometry.

**Figure 4:**
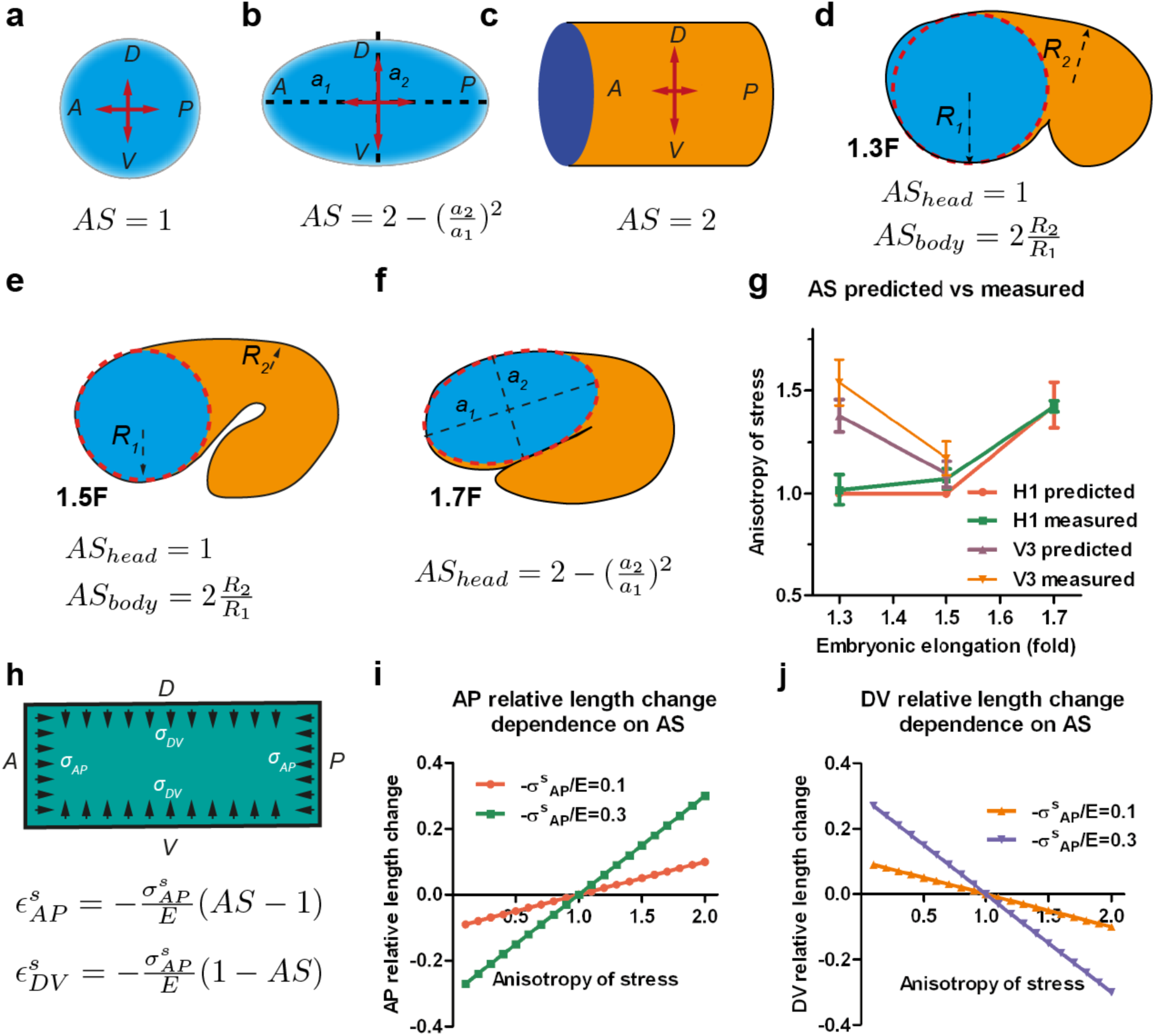
Stress anisotropy induces embryonic morphological changes. (**a, b, c**) Anisotropy of stress (AS) for a sphere, an ellipsoid and a cylinder with DV and AP axis defined in the schemes; (**a, b**) show the middle plane; the major and minor axis of the ellipsoid are called *a_1_* and *a_2_.* (**d, e, f**) The embryo is schematized with a spherical (1.3F and 1.5F stages) or ellipsoidal (1.7F stage) head, and a curved cylindrical body. The AS in the head evolves from 1 (sphere) to that of an ellipsoid, whereas the body AS depends on the ratio of body to head radius (*R_2_/R_1_).* (**g**) Comparison of the predicted AS based on embryo diameter measurements (see figure 3c) and the measured AS obtained from laser ablation experiments (figure 3b). (**h**) Hooke’s law written for an isotropic material like seam cells (supplementary information SI6A); 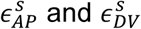 are the relative length changes along the AP and DV directions, respectively. The stress 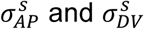 along the AP and DV directions are supposed to be contractile (so negative). E, seam cell Young modulus; A, anterior; P, posterior; D, dorsal; V, ventral. (**i, j**) Dependence of the AP and DV relative length change on the anisotropy of stress for different values of 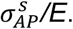

To examine whether the AS can dictate embryonic morphological changes, we related the deformation of the vessel wall with the forces applied using the Hooke’s law (figure 4h, supplementary SI6A) – for instance Hooke’s law states that the one-dimensional deformation of a spring equals to the ratio of the applied force to the spring stiffness. Similarly in a two-dimensional system and for an isotropic material, the deformation is proportional to the mechanical stress (forces) and inversely proportional to the Young modulus (stiffness) along the different loading directions (figure 4h). The resulting equations, which assume that seam cells have an isotropic cortex and are subjected to contractile stress, correctly predict that the seam cell dimension increases along the AP axis *(ε_AP_*) with the AS (figure 4h-j), and decreases along the DV axis *(ε_DV_*). Indeed, consistent with the equations, the head evolves from a sphere to an ellipsoid between the 1.5F to the 1.7F stages as the AS becomes greater than 1 (figure 3b-c).

In conclusion, our experimental and modelling data show that the AS induces morphological changes occurring in embryonic seam cells and provide a basis to understand how the embryo elongates from a mechanical standpoint.

### Stiffness anisotropy-based elongation of the DV epidermis

As shown in figure 3e, the head diameter reduction primarily involves changes of the circumferential width in the DV epidermis. Since the RhoGAP RGA-2 maintains myosin II activation in these cells at a low level (Diogon, 2007), actomyosin contractility in DV cells cannot account for such changes. However, DV epidermal cells have circumferentially-oriented actin bundles, in contrast to seam cells (figure 1b-d), which based on recent observation could affect cell stiffness (Calzado-Martin, 2016, Salker, 2016). We thus hypothesized that the circumferential polarized actin distribution in DV epidermal cells could induce higher stiffness in that direction and thereby influences their deformation. To establish whether it is the case, we investigated both stress and stiffness distribution in the epidermal cells dorsal and ventral to the H1 seam cell using laser nano-ablation (figure 5a). Since these cells are the precursors of the HYP7 syncytium, we will denote them HYP7 henceforth.

**Figure 5:**
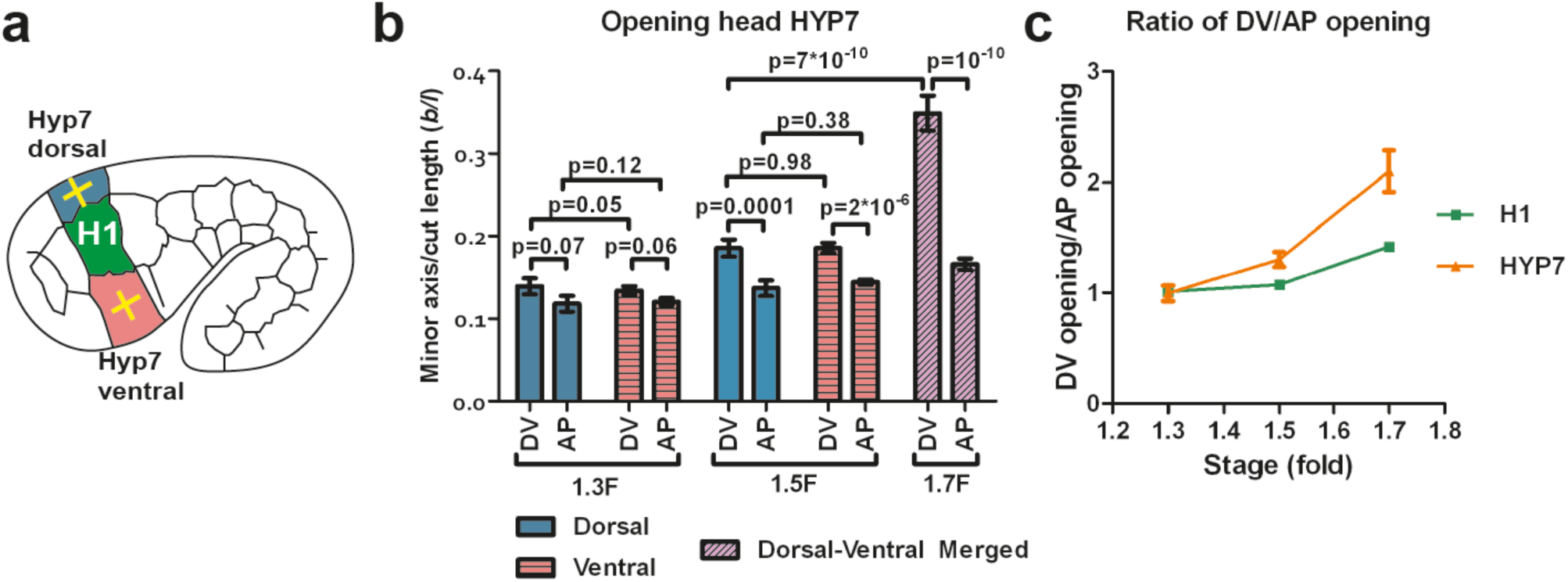
The dorso-ventral epidermis behaves differently than the H1 seam cell in ablation experiments. (**a**) Scheme showing laser ablation experiments in the epithelial cell HYP7 dorsal and ventral to H1, yellow crosses show cut directions. (**b**) Cut opening in the DV and AP directions measured in HYP7 between the 1.3F and 1.7F stages (see figure 2b). p-value of two-tailed t-test is reported. (**c**) Comparison of the DV/AP opening ratio in the seam cell H1 and head HYP7 cell. The data was derived from 3b and 5b. To simplify, we will call the cells participating in the future HYP7 syncytium the HYP7 cells. The number of embryos used for ablation is given in supplementary table 1.

We found that, in the HYP7 cell, the opening in the DV direction was larger than in the AP direction (figure 5b; dorsal and ventral cells behaved similarly after laser cutting), at the 1.5F and 1.7F stages, similarly to the H1 cell (figure 3b). However, the ratio of DV/AP opening in HYP7 was more important than in H1 (figure 5c). Assuming that the HYP7 cell cortex has isotropic material properties like that of H1, our model (figure 4; supplementary SI5) would predict that the DV/AP opening ratio in HYP7 depends only on the head axisymmetric shape and is equal to that of H1, and would thus contradicts our observations. Hence, this *reductio ad absurdum* argument suggests that the HYP7 cell has anisotropic cortical material properties indeed.

To model the DV epidermal cell deformation, we examined two classes of anisotropic stiffness material previously described: orthotropic materials such as bones (Miller, 2002, Helwig, 2009), and fiber-reinforced materials such as arteries (Gasser, 2006), articular cartilage (Federico and Gasser, 2010) or fibrous connective tissues (Ben Amar, 2015). Orthotropic materials have different stiffnesses along orthogonal directions, and thus respond differently to the same stress magnitude along these directions. Fiber-reinforced materials also have different stiffnesses in the directions along and transverse to the fibers; in addition, such materials can respond differently to extensive or compressive stress (Bert, 1977). To define which model best applies to the DV epidermis, we used continuum linear elastic analysis (Muskkhelishvili, 1975, Suo, 1990, Theocaris, 1986, Yoffe, 1951) (supplementary SI7) to interpret the laser cutting data on the DV epidermis. We discarded the orthotropic model, as it did not adequately describe our data (supplementary SI8), and focused on the fiber-reinforced plane model, which better accounts for the presence of well aligned actin fibers in DV cells.

In a fiber-reinforced material composed of a matrix superimposed with fibers, the contribution of the fibers to the stiffness of the material depends on their orientation. In the direction parallel to the fibers, the Young modulus is much increased, whereas in the direction perpendicular to the fibers this contribution is small. According to our modelling, the Young modulus along the fiber direction, increases linearly with a factor *K* related to the fiber stiffness and density; whereas the stiffness along the direction transverse to the fibers varies as a hyperbolic function of *K* and reaches a plateau (supplementary SI7). For fiber-reinforcement in the DV direction, the change in Young modulus along the DV and AP directions predicted by modelling is given in figure 6ab. Cuts perpendicular to the fibers opened similarly to an isotropic material with the matrix Young modulus, because they locally destroyed the fibers (figure 6c; see equation 1 above). By contrast, cuts along the fibers opened with an equilibrium value that depends on the fiber stiffness and distribution through the factor *K* defined above (figure 6d, supplementary SI7).

**Figure 6:**
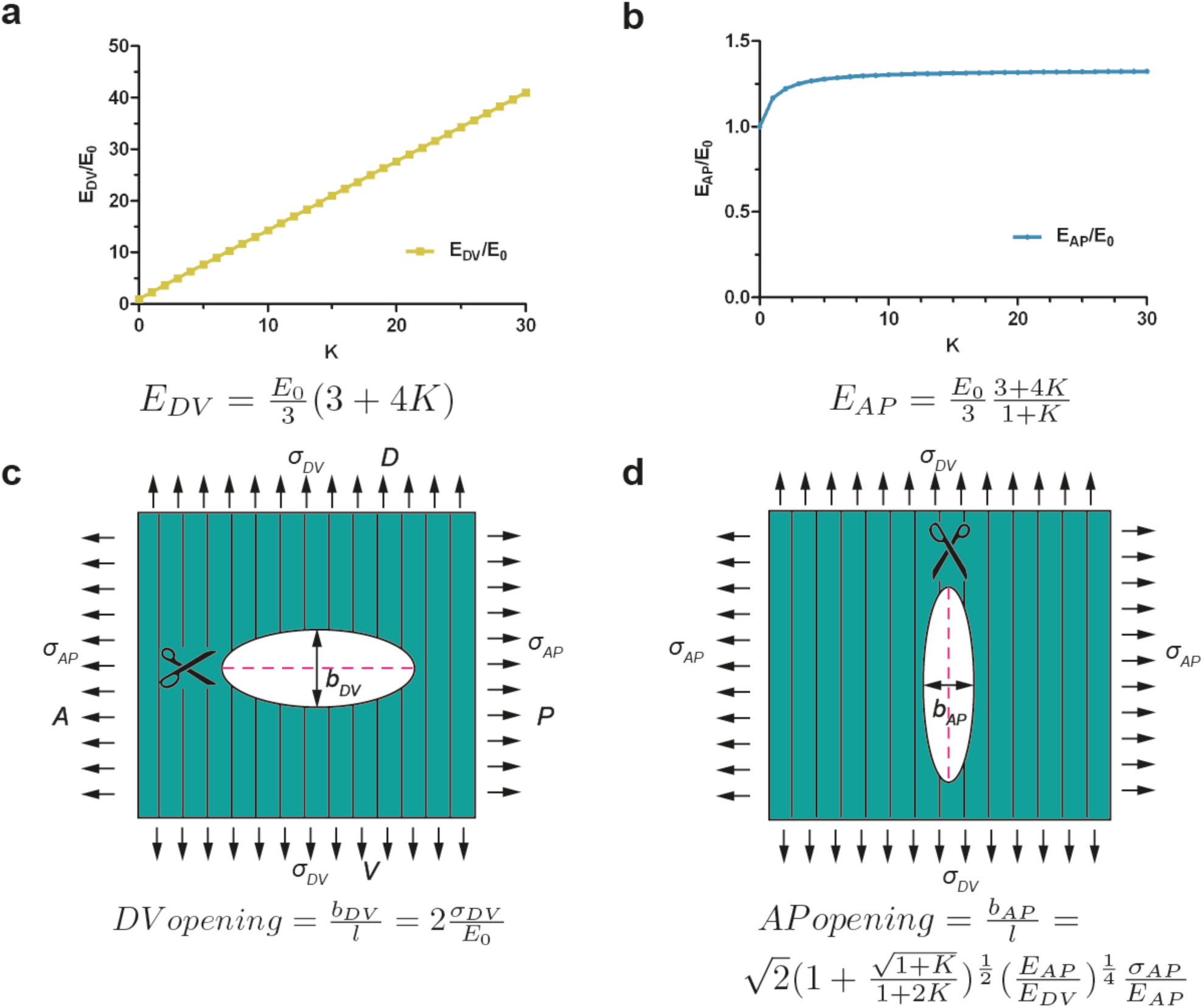
Model of cut opening for a fiber-reinforced material. (**a**) Considering a composite material with fiber reinforcement along the DV direction, the ratio of the Young modulus of the composite material along the DV direction, *E*_DV_, to the Young modulus of the material without fibers (matrix), *E*_0_, depends linearly on a factor *K* (see Supplementary SI7); *K* is related to fiber density and stiffness. (**b**) The ratio of the Young modulus along the AP direction, *E_AP_*, to the matrix Young modulus, *E*_0_, varies little with *K*. (**c**) The opening to the cuts perpendicular to the fibers is similar to an isotropic plane response, and depends on the ratio of DV stress to the matrix Young modulus *σ_DV_*/*E*_0_. (**d**) The opening to the cuts parallel to the fibers depends on the factor *K*, the ratio of AP/DV stiffness, *E_AP_*/*E_DV_*, and the ratio of stress over Young modulus in the AP direction, *σ_AP_*/*E_AP_*.

Since the H1 seam and the head HYP7 cells are adjacent along the circumference (figure 3d), they should be under the same DV stress due to tension continuity across cell-cell junctions. According to equation (1), if the stress in two cells is the same their opening should vary inversely with their respective Young moduli. Since the DV opening of HYP7 was about 1.5 times smaller than that of H1 (figure 7a), we infer that the Young modulus of the HYP7 matrix without fibers was about 1.5 times stiffer than that of H1 (supplementary SI9), suggesting that these cells have distinct material properties. Comparing the DV and AP opening for the HYP7 cell, we found that the factor *K* increased during early elongation (figure 7b; supplementary SI10). More importantly, the calculated ratio of DV/AP Young moduli also increased, and was greater than the DV/AP stress anisotropy (figure 7c).

**Figure 7:**
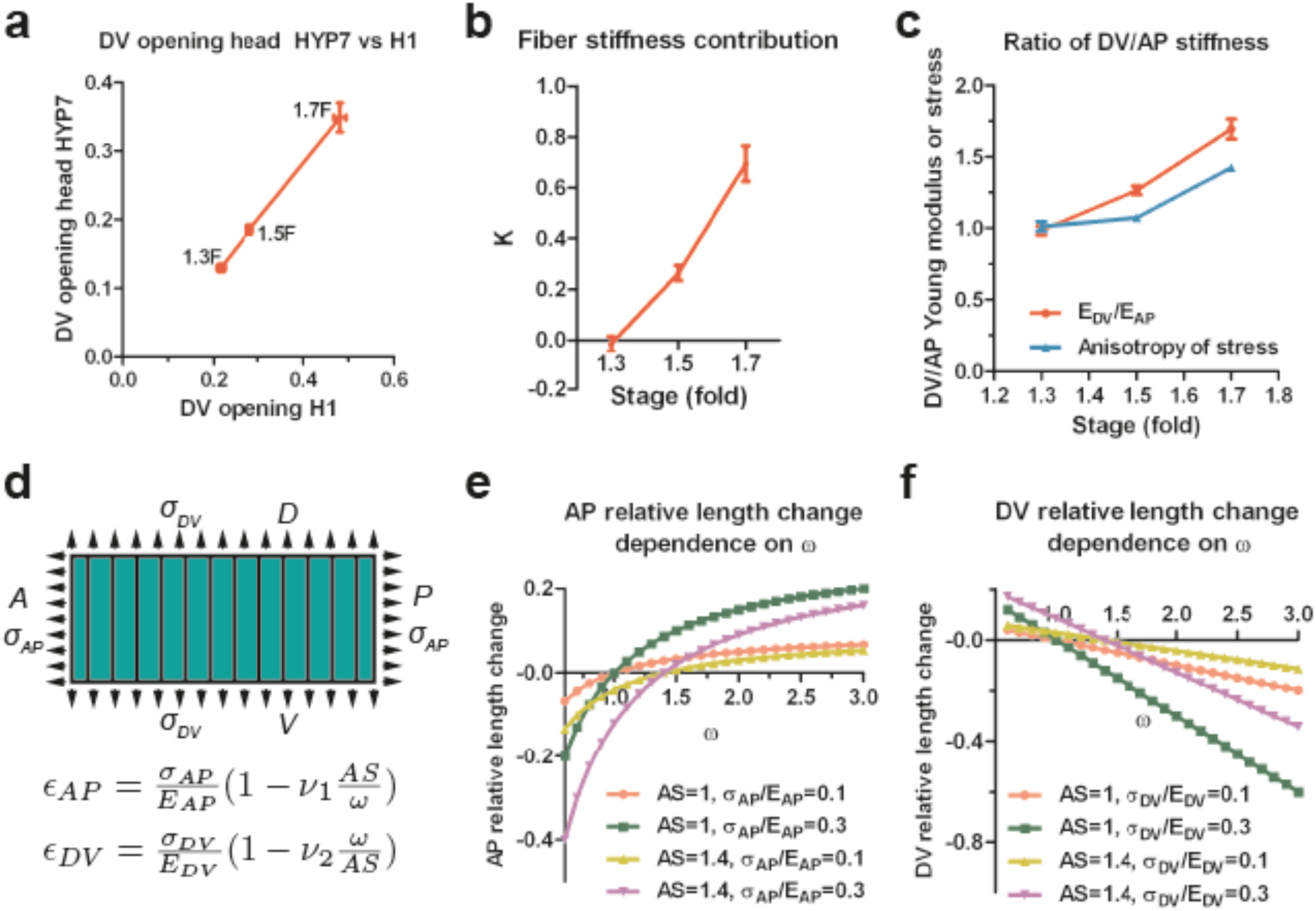
The anisotropy of stiffness in the HYP7 cell helps the embryo to elongate. (**a**) The cut opening in HYP7 is linearly related on the cut opening in H1. The slope of the linear regression gives the ratio of the HYP7 matrix without fibers to H1 Young moduli. (**b**) The factor *K* increases during elongation from the 1.3F to 1.7F stages. (**c**). The ratio of DV/AP stiffness increases and is greater than the AS during early elongation. The data was derived from 3c and SI10. (**d**) Hooke’s law written for a fiber-reinforced material such as DV cells (supplementary information SI6B). *ε_AP_* and *ε_DV_* are the relative length change along the AP and DV directions, respectively. The stress *σ_AP_* and *σ_DV_* along the AP and DV directions are supposed to be tensile (so positive). *E_AP_* and *E_DV_* are the Young moduli in the DV cells along the AP and DV direction, respectively. *v_1_* and *v_2_* are Poisson’s ratios in DV cells; *ω* is the *E_DV_*/*E_AP_* ratio. A, anterior; P, posterior; D, dorsal; V, ventral. (e) Dependence of the AP relative length change *ε_AP_* on the ratio of DV/AP stiffness ω for different value of AS and *σ_AP_*/*E_AP_*; the *v_1_* value is taken to be 1. (**f**) Dependence of the DV relative length change *ε_DV_* on the ratio of DV/AP stiffness *ω* for different value of AS and *σ_DV_*/*E_DV_*; the *v_2_* value is taken to be 1.

To understand how a change in stiffness affects head HYP7 deformation, we wrote again Hooke’s law for these cells (supplementary SI6B; figure 7d). Since myosin II activity in DV cells is low, their cortex should be exposed to tensile stress induced by actomyosin contractility in the seam cells. The cell length along the AP direction increased when the stiffness anisotropy (DV/AP stiffness ratio) increased (figure 7d-e), whereas the trend was opposite in the DV direction (figure 7d,f). Thus the stiffness anisotropy helps the HYP7 cell extend along the AP direction and shrink along the DV direction. Interestingly, the equations predict that increasing stress anisotropy has an opposite effect on HYP7 cell deformation, since it prevents them from extending antero-posteriorly (figure 7e). Altogether, our model strongly suggests that when the DV/AP stiffness anisotropy increases and is higher than the DV/AP stress anisotropy, as observed in the head HYP7 (figure 7c), elongation along the AP direction is favoured. Furthermore, our data highlight that the distinct mechanical properties of cells composing a complex tissue enables its morphogenesis and does not require all cells to be contractile.

### The stress and stiffness anisotropies correlate with actin arrangement

Since myosin II is not polarized (figure 1e-f), to find out if actin distribution accounts for the stress and stiffness anisotropies, we carried out an analysis of actin filament alignment in the seam cells H1 and V3, as well as in the head HYP7 cell. We found that the polarization of actin filaments in seam cells correlated with the observed pattern of stress anisotropy. Indeed, in H1 at the 1.3F stage, actin filaments had a nearly isotropic angular distribution correlating with the isotropic stress (figure 8a, figure 3b), whereas they became increasingly aligned along the DV direction from the 1.3F to 1.7F stages (figure 8a) which mirrors the increase of stress anisotropy (figure 3b). Likewise, actin alignment decreased along the DV direction in V3 from the 1.3F to the 1.5F stage, in parallel to the decrease of stress anisotropy between those stages (figures 8b and 3b). The changes in H1 (figure 8c) were statistically significant, whereas it is not the case in V3 (figure 8d). In the HYP7 cell, actin filaments already acquired a preferential DV alignment at the 1.3F stage (figure 8c), but became increasingly organized along the DV direction as the embryo elongated to the 2F stage, with a highly significant difference between the 1.5F and 1.7F stages (figure 8c and 8f). These changes correlated with the increase of stiffness anisotropy observed in the HYP7 cell (figure 7c).

**Figure 8:**
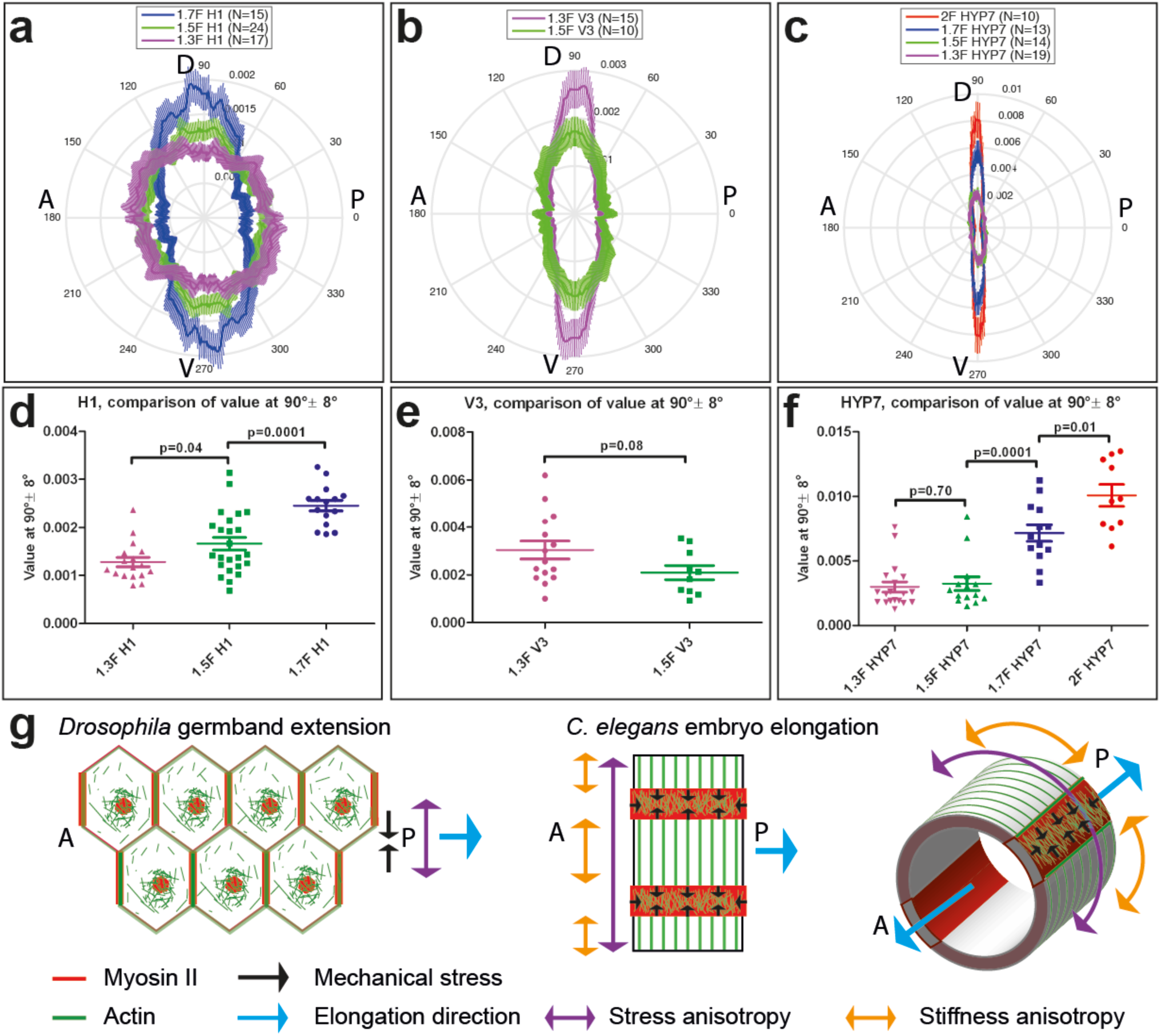
Actin filament organization correlated with stress and stiffness anisotropy pattern. (**a-c**) Angle distribution of actin filaments in the seam cell H1 (a), seam V3 (b) and the HYP7 cell (c), at different elongation stages. D, dorsal; V, ventral; A, anterior; P, posterior. 90° correspond to DV direction. (**d, e, f**) Comparison of the peak values at 90°± 8° (DV direction) of angular distribution showed in (**a, b, c**) respectively. p-value of two-tailed t-test is reported. (**g**) (**left**) The anisotropy of mechanical stress generated by the polarized actomyosin network and medial myosin pulses promote *Drosophila* germband extension; (**right**) The interplay of stress anisotropy (generated in seam cells - red) and stiffness anisotropy (DV cells – white) promote *C. elegans* embryo elongation. Note that, while myosin II does not display a polarized distribution within individual *C. elegans* epidermal cells like in *Drosophila* germband epithelial cells, its enrichment in seam cells along the circumference is reminiscent of the localized myosin II enrichment at vertical junctions in *Drosophila.* A, anterior; P, posterior.

We have attempted to functionally test how actin organization could affect stress and stiffness by manipulating actin polymerization through two different strategies to express cofilin during early elongation. However, we could not obtain meaningful results. Altogether, we conclude that the pattern of actin distribution showed a good correlation with the observed stress and stiffness anisotropy. It will remain important to define the mechanisms bringing changes in actin distribution, and ultimately whether it is a cause or a consequence of anisotropy.

## Discussion

Classical experiments in embryology have outlined how the juxtaposition of cells with different properties is critical to power important morphogenetic movements, such as *Xenopus* gastrulation (Hardin and Keller, 1988, Keller and Winklbauer, 1992). In this work, we have dissected the mechanical contributions of the different epidermal cells driving *C. elegans* embryonic morphogenesis at the single cell resolution, highlighting the importance of juxtaposition of cells with different properties. Combining laser nano-ablation and continuum mechanics modelling, we first highlight the importance of stress anisotropy in the seam cells. Second, we emphasize that stiffness anisotropy is equally important for embryonic elongation but matters in another epidermal cell type, the DV cells. Thereby, we reveal the critical role of tissue material properties in morphogenesis.

Many studies analysing morphogenetic processes have focused on 2D epithelial sheets such as the *Drosophila* mesoderm (Martin, 2009), germband (Rauzi, 2008, Rauzi, 2010, Blankenship, 2006, Fernandez-Gonzalez and Zallen, 2011), amnioserosa (Solon, 2009, Gorfinkiel, 2009), wing and thorax (Aigouy, 2010, Bosveld, 2012), or the zebrafish enveloping cell layer (Behrndt, 2012) during embryonic development. They have revealed the role of contractile actomyosin pulses and planar polarity in coordinating events over long distances. The *C. elegans* embryonic elongation is distinct from those situations since it does not involve myosin II polarized distribution nor actomyosin pulses. Interestingly, this process still requires stress anisotropy, outlining that it can be reached by different means. We suggest that several factors contribute to establish stress anisotropy in *C. elegans.* First, the actin network displayed a more polarized dorso-ventral distribution in seam cells when the stress anisotropy was higher, which should increase the stress in that direction. Second, akin to a planar polarized distribution, myosin II activity displays an asymmetric distribution along the embryo circumference in cells with different material properties (figure 8g). Intriguingly, tissue culture cells can sense the spatial stiffness distribution (Walcott and Sun, 2010, Fouchard, 2011, Trichet, 2012, Lange and Fabry, 2013), raising the possibility that seam cells sense the higher circumferential stiffness and respond with higher DV-oriented stress through the mechanosensitive adherens junctions (le Duc, 2010, Yonemura, 2010). Third, we found that the spectrin cytoskeleton matters to reach normal levels of stress magnitude and anisotropy. Spectrin is known to impinge on actin filament alignment and continuity in DV cells (Praitis, 2005, Norman and Moerman, 2002) and could thus affect DV stiffness anisotropy by reducing the level of actin fiber alignment. Finally, although myosin II activity is low in DV cells (Diogon, 2007), the remaining activity might create some DV-oriented stress feeding back on seam cells.

By modelling the DV cells as a fiber-reinforced material, we reveal how the polarized cytoskeleton in DV cells increases their stiffness to orient the extension in the AP direction, acting like a ‘molecular corset’. Related ‘molecular corsets’ have been described and proposed to drive axis elongation in other systems (Wainwright, 1988). In *Drosophila*, a network of extracellular matrix fibrils was proposed to help elongate developing eggs (Haigo and Bilder, 2011). In plant cells, the orientation of cellulose microfibrils determines the axis of maximal expansion. In the latter, stiffness anisotropy also helps overcome stress anisotropy (Green, 1962, Baskin, 2005). Importantly, *C. elegans* embryos reduce their circumference during elongation, while *Drosophila* eggs and plants increase it. It suggests that to conserve the actin reinforcement properties when the diameter decreases, *C. elegans* DV epidermal cells should have a mechanism to actively shorten the actin bundles, as observed in a biomimetic *in vitro* system (Murrell and Gardel, 2012).

Our experimental data were consistent with the predictions from Hooke’s law. They prove that the actomyosin cortex preferentially squeezes the embryo circumferentially, and that the stress anisotropy is tightly linked to the geometry of the embryo. By quantitatively assessing the contribution of stiffness anisotropy in tissue elongation, we have emphasized its importance relative to the more established role of stress anisotropy. The precise relationship between both anisotropies remains to be investigated. Thus, the juxtaposition of cells with different "physical phenotypes”, seam epidermis expressing stress anisotropy and DV epidermal cell showing stiffness anisotropy, powers *C. elegans* elongation, as previously suggested in chicken limb bud outgrowth (Damon, 2008) or chick intestinal looping (Savin, 2011). We did not mention other potential stress bearing components, like microtubules and the embryonic sheath (Priess and Hirsh, 1986), since the former mainly serves to enable protein transport (Quintin, 2016) whereas the function of the later will be the focus of an upcoming work.

In conclusion, our work highlights that tissue elongation relies on two fundamental physical quantities (mechanical stress and tissue stiffness), and provide the most advanced mesoscopic understanding to date of the mechanics at work during the first steps of *C. elegans* embryonic elongation.

## Methods

### *C. elegans* alleles and strains

Bristol N2 was used as the wild-type (WT) strain and animals were maintained as described in Brenner (Haigo and Bilder, 2011). The strain ML1540: *mcIs50[lin-26p::vab-10(abd)::gfp; myo-2p::gfp]* LGI carrying the actin-binding domain (ABD) of the protein VAB-10 under the epidermal promoter *lin-26* was described elsewhere (Green, 1962, Baskin, 2005). The endogenous NMY-1::GFP reporter strain was built by CRISPR knock-in (ML2540: *nmy-1(mc82)[nmy-1::gfp]* LGX; the NMY-2::GFP reporter strain LP162 *nmy-2(cp13)[nmy-2::gfp* + *LoxP]* LGI was a generous gift from Daniel Dickinson.

For parallel calcium and actin imaging during ablation, we used the strain ML2142: *mcIs43 [lin26p:: vab-10::mCherry; myo-2p::gfp]; juIs307[dpy-7p::GCaMP3])* carrying a calcium sensor under the epidermal promoter *dpy-7p* and mCherry-labeled VAB-10(ABD) under the *lin-26* promoter. The thermosensitive Rho kinase mutation *let-502(sb118ts)* was crossed with ML1540 to give the strain ML2216: *let-502(sb118ts); mcIs50[lin-26p::vab-10(abd)::gfp; myo-2p::gfp]* LGI.

For determining morphological changes, we used the strain ML2386: *mcIs50[lin-26p::vab-10(abd)::gfp; myo-2p::gfp] I; xnIs97[hmr-1::gfp] III*) expressing both a junctional marker (HMR-1/E-cadherin) and an actin marker (VAB-10(ABD)).

For actin alignment analysis we used the ML1966 *unc-119(ed3) mcIs67[dpy7p::LifeAct::GFP; unc-119(*+)] strain, expressing the actin reporter LIFEACT under the *dpy-7* epidermal promoter.

### RNA interference

RNAi experiments were done using injection of double-stranded RNA synthesized from PCR-amplified genomic fragments using a T3 or T7 mMESSAGE mMACHINE Kit (Ambion, Austin, TX, USA). The embryos were analyzed from 24h to 48h post-injection.

### Time-lapse analysis and morphological change quantification

Freshly laid embryos or embryos from dissected hermaphrodites were mounted on 5% agarose pads in M9 buffer and the coverslip was sealed with paraffin oil. DIC time-lapse movies were recorded at 20°C using a Leica DM6000 upright microscope with a 40X oil immersion objective. For each embryo, a Z-stack of 7-8 focal planes with 4 μm step size was acquired. The length of embryos was estimated by tracing the embryo body axis (through the middle of the embryo). Fluorescence time-lapse movies were recorded at 20°C using a spinning-disk Zeiss microscope Axio Observer.Z1 using a 63X oil immersion objective. Other fluorescence images were acquired with the same microscope using a 100X oil immersion objective. To determine the morphological changes of the embryo, sections of the embryo imaged with junctional and actin markers at the level of H1, V3 and V6 were reconstructed to determine the radius, seam and DV cell width along the circumferential direction. All images were analysed using the ImageJ (FiJi) software (NIH, Bethesda, Maryland, USA; http://rsb.info.nih.gov/ij/) and MATLAB R2014b (The MathWorks Inc., Natick, MA).

### Actin alignment analysis

Z-stack images of LIFEACT::GFP fluorescence expression in the epidermis were acquired using a confocal Leica SP5 microscope with a 63X oil immersion objective and zoom factor 8. We used a step size of 0.08 μm a pinhole opening of 0.6 Airy Unit and projected 2 μm around the actin cortex. The embryos were rotated on the scan field to have the same antero-posterior orientation. The acquired images were deconvoluted using the Huygens Essential software from Scientific Volume Imaging (Hilversum, Netherlands). We chose a region of interest (ROI) of 4×4 μm^2^ within the seam cell H1 or dorso-ventral epidermal cell HYP7, and of 3×3 μm^2^ within the seam cell V3 to perform Fast Fourier Transform (FFT). We used a high-pass filter to remove the low frequencies then did inverse FFT. We found that the high pass filter removed changes in intensity due to unequal labelling or out of focus signals but retain the actin texture. Finally we use an ImageJ plugin “Spectral Texture Analysis” written by Julien Pontabry to derive the angle distribution of actin texture. This plugin performed the Fast Fourier Transform (FFT) of the given ROI and computing coefficients in Fourier space, such as the angle distribution of the given structure, as detailed in Gonzalez, R.C. & Woods, R.E. (2008). Digital image processing, Nueva Jersey, chapter 11, section 3.3 (Murrell and Gardel, 2012).

### Embryo staging for ablation

For ablations, we compared embryos of the same developmental timing. To do so, we recorded the elongation curve of different genetics background (Figure 3-Figure supplement 1) and took embryos at the corresponding developmental time from the beginning of elongation. Thus, *unc-112(RNAi)* embryos elongating up to 1.7F similarly to WT have the same length as WT at 1.7F stage. In contrast, *spc-1(RNAi)* embryos elongated slower than WT (figure S4) and thus at a time corresponding to 1.7F in a control embryos were shorter than wild-type embryos 1.7F stage. For comparison between *let-502(sb118ts)* embryos, measurements were carried out at 25.5°C and the embryos were taken when muscles start to twitch (around 1.5F in control embryos).

### Laser ablation

Laser ablation was performed using a Leica TCS SP8 Confocal Laser Scanning microscope, with a femtosecond near-infrared Coherent Chameleon Vision II, Ti:Sapphire 680-1080 nm laser, 80 MHz. To make a line cut, a region of interest with a length varying from 3 to 6 μm and a width of 0.08 μm (1 pixel width) was drawn. We used a laser wavelength varying from 800 nm to 900 nm, which gave similar ablation responses. The laser power was tuned before each imaging section to obtain local disruption of the cortex response (>80% of the cases, visible opening, no actin accumulation around cell borders in the repair process and the ablated embryos developed normally). Typically the power of the laser was 2000 mW, and we used 50% power at 100% gain. Wounding response (actin accumulation around cell borders in the repair process, embryo died afterwards) was rarely observed at the power used for local disruption, but more often when the power was increased to 60-65%. The first time point was recorded 1.44 s after cutting, which corresponded to the time needed to reset the microscope from a two-photon to a regular imaging configuration. The image scanning time recorded by the software was usually less than 400 ms, so the total exposure time of the chosen ROI to multiphoton laser was less than 1 ms. The cuts were oriented either in the antero-posterior (AP) or dorso-ventral (DV) directions relative to the global orientation of the embryos. After ablation, the embryos were monitored to see if they continued to develop normally or they expressed the desired phenotype. More precisely, we verified whether embryos ablated at 1.3F and 1.5F developed past the 2F stage, embryos ablated at 1.7F developed past the 2.5F stage, *unc-112(RNAi)* embryos and *spc-1(RNAi)* embryos arrested at 2F stage.

### Laser ablation image processing and data analysis

The shape of the cut opening was detected using the Active Contour plugin ABsnake (Brenner, 1974). A starting ROI was drawn around the opening as the initiation ROI for ABsnake. After running the plugin, the results were checked and corrected for detection errors. The detected shape was fitted with an ellipse to derive the minor axis, major axis and the angle formed by the major axis with the initial cut direction. The average opening of the five last time points before the repair process began (figure S2, from around 8 to 10 s after cutting) was taken as the opening at equilibrium. The standard error of the mean was shown.

The curve fit was performed on the average value of the cut opening (defined as the minor axis/initial cut length) using GraphPad Prism 5.00 (San Diego, California, USA) and the equation of one-phase association:

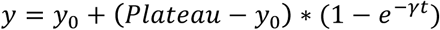

where y_0_ is the initial width of the cut opening, *Plateau* is the minor axis of the opening at equilibrium and γ is the relaxation rate. The standard error of the mean given by the software was shown.

### Statistical analysis

The two-tailed t-test was performed on the average of the last five time points (from about 8 s to 10 s) of the cut opening using MATLAB R2014b (The MathWorks Inc., Natick, MA). Z-test was performed using QuickCalcs of GraphPad Prism (San Diego, California, USA) to compare the anisotropy of stress (AS), the relaxation half-time and the initial recoil speed of the cut opening.

## Acknowledgments

The authors thank Demet Kirmizibayrak and Marcel Boeglin for technical assistance, Pierre-François Lenne, Flora Llense, Teresa Ferraro, François Robin, Sylvie Schneider-Maunoury and Raphaёl Voituriez for critical comments on the manuscript.

This work was supported by an European Research Council grant to ML (grant #294744), and by institutional funds from the Centre National de la Recherche Scientifique (CNRS), University of Strasbourg and University Pierre et Marie Curie (UPMC), the grant ANR-10-LABX-0030-INRT which is a French State fund managed by the Agence Nationale de la Recherche under the frame programme Investissements d’Avenir labelled ANR-10-IDEX-0002-02 to the IGBMC, and by installation grants from the CNRS and UPMC to ML. MBA is supported in part by Institut Universitaire de France. Some strains were obtained from the *Caenorhabditis* Genetics Center CGC (funded by the NIH Office of Research Infrastructure Programs P40 OD010440). Confocal work was carried out at the Institute of Biology Paris Seine Imaging facility that is significantly supported by the “Conseil Regional Ile-de-France”, the French national research council (CNRS and Sorbonne University, UPMC Univ Paris 06.

## Authors’ contributions

T.K.T.V-B and M.L designed the project, the approach and wrote the manuscript. T.K.T.V-B performed all experiments, proposed the calculations in SI2-SI6 and SI8-SI10. M.B.A proposed the modelling in SI7. J.P wrote the "Spectral Texture Analysis” plugin.

## Supplementary movie legends

**Supplementary movie 1:** local disruption of actin cortex with laser ablation, visualized with the actin marker (ABD::mCHERRY) expressed under the epidermal promoter *lin-26.* 0 s time corresponds to first picture after laser cut. Yellow line shows the cut region.

**Supplementary movie 2:** local disruption of actin cortex with laser ablation does not induce noticeable change in calcium level. The calcium sensor GCaMP3 was expressed under the epidermal promoter *dpy-7* (strong expression in the dorso-ventral cells). 0 s time corresponds to first picture after laser cut. Yellow line shows the cut region. It is the same embryo as shown in movie 1.

**Supplementary movie 3:** wound healing response after laser ablation visualized with the actin marker ABD::mCHERRY expressed under the epidermal promoter *lin-26.* 0 s time corresponds to first picture after laser cut. Yellow line shows the cut region.

**Supplementary movie 4:** calcium wave propagation in wound healing response after laser ablation. The calcium sensor GCaMP3 was expressed under the epidermal *dpy-7* promoter (strong expression in the dorso-ventral cells). 0 s time corresponds to first picture after laser cut. Yellow line shows the cut region. It is the same embryo as shown in movie 3.

